# BIQ: A method for searching circular RNAs in transcriptome databases by indexing backsplice junctions

**DOI:** 10.1101/556993

**Authors:** Peter Menzel, Irmtraud M Meyer

## Abstract

Circular RNAs (circRNAs) are a class of RNA transcripts that originate from non-canonical splicing events and are characterized by a backsplice junction connecting the 3’ splice site to an upstream 5’ splice site. Here, we present the program *BIQ* for indexing and querying transcriptome sequencing datasets for backsplice junctions. BIQ can be used for instantaneously querying all indexed transcriptomes for occurrence and abundance of reads overlapping the backsplice junction of a particular circular RNA, which can help in the functional characterization of known and novel circular RNAs. BIQ is free software and available at https://github.com/pmenzel/biq.

## Introduction

Circular RNAs (circRNAs) are a type of transcripts that has only recently been found to be widely abundant in metazoan cells (1, 2). They originate from non-canonical splicing events in which the 3’-end of an RNA molecule is spliced to its 5’-end, also called *backsplicing* (3), creating splice junctions that are denoted as backsplice junctions (BSJ). The majority of circular RNAs are the product of backsplicing an exon’s 3’-end to its 5’-end or to the 5’-end of an upstream exon, possibly retaining inner introns and exons (4–6). This type of backsplicing is facilitated by the sequence content of the flanking introns (7, 8). Similar to regular splicing, one gene can give to rise to multiple distinct circular RNA transcripts, which can be distinguished by their backsplice junctions (9). For example, at least five circular RNA isoforms have been found in the human gene *HIPK3*, of which the one containing only the second exon, *circHIPK3*, is the most abundant (10).

Analysis of transcriptomes from total RNA libraries of multiple model organisms demonstrates a large variety of circular RNAs among different cell lines and tissue types (2, 3, 6). One hallmark of circular RNAs is their enrichment in the mammalian brain and nervous system (11, 12). Similarly, circular RNA abundance increases throughout the development of *Drosophila melanogaster* and the highest enrichment is observed in the nervous system (13). Most circular RNAs are exported to the cytoplasm and, presumably, due to their resistance to degradation by exonucleases, have extended lifetimes compared with linear transcripts (1, 7). Besides these global observations about circular RNA abundance and diversity, only few circular RNAs have been characterized by their putative molecular functions and their involvement in biological processes. For example, cytoplasmic circular RNAs could act as "sponges" for microRNAs, such as the highly abundant human circular RNA *CDR1as*, which contains more than 70 binding sites for *mir-7* (4, 14), or *circHIPK3*, which contains binding sites for several microRNAs including *mir-124* (10). The latter example particularly suggests that *circHIPK3* is a critical functional product of the *HIPK3* gene, and is involved in neural gene regulatory networks, in particular impacting cell proliferation. Therefore, by modulating gene expression, circular RNAs may also be involved in tumorigenesis and could serve as diagnostic biomarkers or therapeutic targets (15, 16).

Baélicing can be detected by transcriptomic analysis, for example using short read RNA-Sequencing from libraries created from the total RNA population in a sample. Before library preparation, the RNA can also be enriched for circular RNAs by applying *RNase R* treatment for digesting linear isoforms (7). Several software packages for detecting circular RNAs in RNA-Seq data have been developed in recent years (9, 17). The underlying principle of these programs is the alignment of sequencing reads to a reference genome by employing a read mapping program that is able to split-map reads, such as STAR (18) or BWA-MEM (19). Split-mapping does not require a contiguous alignment of the read to the reference genome, but allows for different sections of the read to be mapped "out-of-order". Using such chimeric alignments, backsplice junctions can be identified as alignments in which a read’s 3’-end is aligned before its 5’-end, which are further filtered by coverage and presence of flanking splicing signals.

One fundamental approach for deciphering the biological processes in which a particular circular RNA may be involved is to measure its abundance across different tissues or cell types and under varying conditions using RNASequencing. Given the large number of already available RNA-Seq datasets in public databases, such as the NCBI sequence read archive (SRA), it is desirable to be able to quickly lookup a circular RNA’s abundance among available transcriptome datasets from a variety of sources.

Here, we present a lightweight and fast method for indexing and querying backsplice junctions in transcriptome datasets, called *BIQ (Backsplice junction Indexing and Query)*. It can be used to index a set of BSJs in a large corpus of transcriptome datasets and query this index for particular backsplice junctions. The analysis of the whole index could also aid in identification of previously unknown circular RNAs, and the quantification of BSJ abundances across all experiments in one index can further aid in generating hypotheses regarding a circular RNA’s putative biological roles.

## Results

### BSJ indexing and querying

The overall workflow of BIQ is outlined in Figure 1. First, BIQ builds an index of *k*mers that span backsplice junctions, which can either be from an enumeration of all possible exon-exon 3’-5’ linkages in a given set of annotated genes and/or from a set of previously detected backsplicing sites. This fixed list of *k*-mers is then used for detecting and counting expressed BSJs in transcriptome datasets, e.g., from the Sequence Read Archive (SRA), and storing them in a database file. This database can then be queried for the backsplice junction, represented by the BSJ-spanning *k*-mer, of a particular circular RNA of interest and BIQ returns the names of the datasets containing at least one sequencing read spanning the query BSJ as well as the *k*-mer abundances and normalized counts. By using a fixed static set of *k*-mers, the database file remains small, even when containing thousands of datasets and the time for querying a single BSJ is near-instantaneous. BIQ uses a *k*-mer length of *k* = 32, comprising 16 nt on each side of the BSJ.

**Fig. 1.**
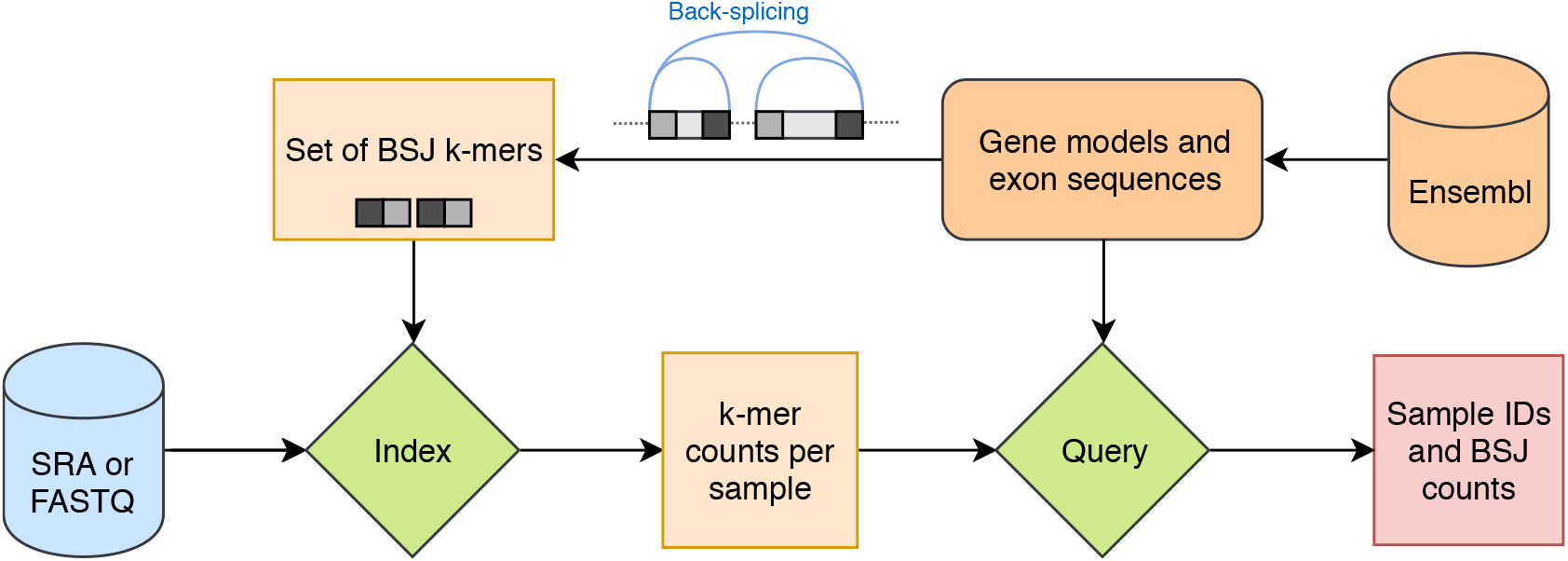
Flow-chart of enumerating, indexing and searching of backsplice-junctions in transcriptome sequencing datasets using BIQ.

### Case study

For illustrating BIQ’s ability to detect circular RNAs, we use two example datasets with total RNA transcriptomes from the fruit fly and human. The first dataset contains 103 short read transcriptomes from *Drosophila melanogaster*, which comprise various parts or tissues of adult flies as well as whole embryo transcriptomes from various developmental stages and five cell lines (13). This particular dataset was previously used by Westholm et al. for detection and quantification of circular RNAs. The second dataset contains total RNA transcriptomes of 202 human samples from various tissues and primary cells as well as from *in vitro* differentiated cells that were sequenced as part of the ENCODE project (20).

First, we create a set of backsplice junction-spanning *k*-mers (BSJ *k*-mers) from the genome annotation. By enumerating all possible 3’-5’ backsplicing junctions from the constituting exons of each gene, we generate ∼180k unique BSJ *k*-mers in the fruit fly and ∼2.9m unique BSJ *k*-mers in human. Next, the BSJ *k*-mers were quantified in all transcriptomes from both datasets and we detected ∼300k occurrences of 9.150 unique BSJ *k*-mers in the fruit fly, and ∼16m occurrences of 253.550 unique BSJ *k*-mers in human (Suppl. Figure 1). In both datasets, the total counts per BSJ *k*-mer follow a distribution in which most BSJ *k*-mers are only found once, whereas only few BSJ *k*-mers are highly abundant (Suppl. Figure 2), which has also been observed in earlier studies (10, 21). Similarly, most BSJ *k*-mers are only found in one sample, whereas few *k*-mers are found in the majority of the samples (Suppl. Figure 3). For example, the three BSJ *k*-mers that occur in most samples in the ENCODE dataset belong to the circular RNAs *circCDYL, circHIPK3*, and *cSMARCA5*, all of which have been associated with cancer progression, such as bladder cancer (22) or hepatocellular carcinoma (23). Despite their rather short length, there is only little overlap between the BSJ *k*-mers and the *k*-mers from regular linear transcripts. Only 8 and 325 BSJ *k*-mers are also found in the annotated linear cDNA sequences from *Drosophila melanogaster* and human, respectively.

Using the BSJ *k*-mer count profiles for each dataset, samples can be compared and clustered just by their circular RNA abundance profiles, and we used UMAP dimension reduction to arrange all samples in two dimensions. In the *Drosophila melanogaster* dataset, we observe that most of the sample types are clustered just by their BSJ profiles (Figure 2a). We can also see that the transcriptomes from cell lines, especially CME and Kc167, are separated from the regular samples. Next, we measured the abundance of BSJ *k*-mers throughout the developmental stages from early embryo to adult (20 day) fly. We observe an increasing number of BSJ *k*-mers throughout embryogenesis (Figure 2b), as was also found by Westholm et al., and, further, the number of distinct BSJ *k*mers in the nervous system also increases throughout the developmental time points and is highest in adult flies (Suppl. Figure 4).

**Fig. 2.**
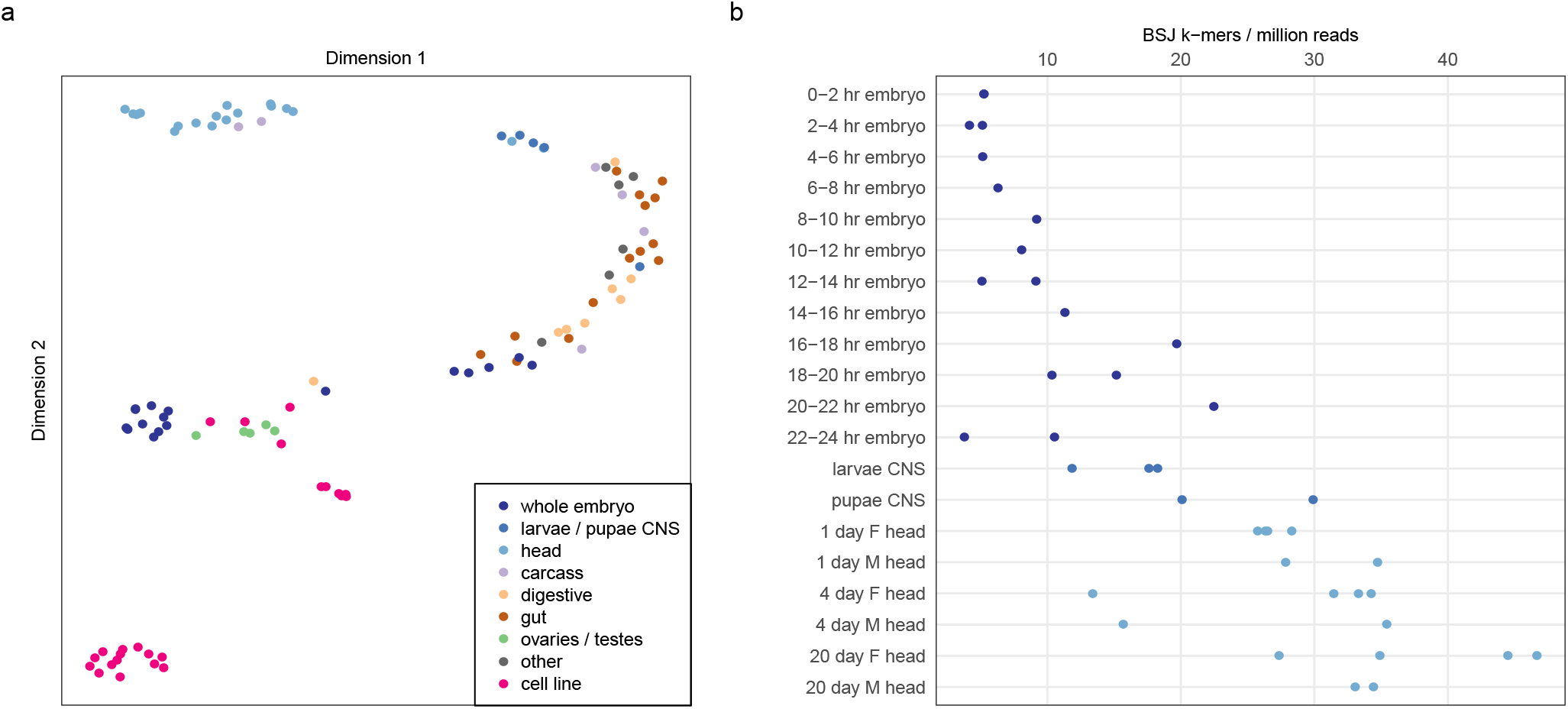
**(a)** Two-dimensional embedding of BSJ profiles of all 103 *Drosophila melanogaster* transcriptomes **(b)** BSJ counts throughout development and in adult fly heads

In the ENCODE dataset, the arrangement of BSJ profiles in two dimensions also shows several distinct clusters belonging to various tissue and cell types, which, however, are more complex compared with the fruit fly dataset, due to the dataset size and overlapping types of source materials. In particular, distinct clusters comprising the central nervous system, thyroid gland, or the colon are visible (Suppl. Figure 5). We also observe, to some degree, a separation of transcriptomes derived from the three types of source materials (primary cells, tissues, and *in vitro* differentiated cells).

## Conclusion

BIQ is a lightweight and fast application for searching circular RNAs in large transcriptome databases by indexing backsplice junction spanning *k*-mers. It contains an easy-to-use graphical user interface, which can be used to browse annotated genes and their exons and query the database for one or multiple backsplice junctions. While its main purpose is fast database indexing and querying, we also showed that BIQ’s *k*-mer based approach can reliably detect circular RNAs and recapitulates the general characteristics of circular RNA abundances when applied to two example datasets from two species.

## Methods

### Implementation

BIQ is implemented as a C++ program that provides a command-line interface both for indexing transcriptomes and querying *k*-mers. The program can read either FASTA/Q files or download datasets from the Sequence Read Archive (SRA) when given a list of accession numbers. For indexing a large collection of transcriptomes, the processing time is mostly governed by file I/O or downloading and extracting SRA files. Regardless, BIQ uses multiple threads for parallel read processing when reading the input data. By default, BIQ reads the gene annotations (in GTF format) and exon sequences from the Ensembl database in order to enumerate all possible exon-exon 3’-5’ combinations to create the initial set of BSJ-spanning *k*-mers. For querying the indexed datasets, BIQ also contains a graphical user interface for viewing genes and their exons sequences in a web browser in order to create a query containing one or multiple BSJ *k*-mers. The search returns a table with experiment IDs and associated read counts as well as counts normalized by library size (in reads per million). The GUI can either be used locally or installed on web server.

### Data analysis

RNA-Sequencing data of all 103 samples from Westholm et al. were downloaded from the Sequence Read Archive (25) and indexed with BIQ using a set of 180.153 unique BSJ *k*-mers (*k*=32), which were derived from enumerating all possible backsplice junctions in protein-coding and lncRNA genes, using the *Drosophila melanogaster* genome annotation from Ensemble. ENCODE samples were selected by filtering the ENCODE experiments by *Assay category* = Transcription, *Assay* = total RNA-seq, and *Organism* = Homo sapiens, using the ENCODE data portal www.encodeproject.org (26), which resulted in 202 experiments comprising 247 SRA files. Again, all possible backsplice junctions were enumerated, resulting in 1.965.630 unique BSJ *k*-mers. After counting the BSJ *k*mers in all samples in each dataset, the BIQ index was exported as a count matrix and *k*-mers that overlap annotated linear transcripts were removed. For visualisation, the count matrix was reduced to two dimensions using UMAP (27). Samples in the Drosophila dataset were manually grouped into broad categories based on their description (see Table S1 in (13)), and samples from ENCODE were grouped by system type based on their biosample ontology ID.

Both example datasets can be queried using the GUI at https://pmenzel.github.io/biq/ and R scripts for recreating the figures are available at https://github.com/pmenzel/biq-manuscript-data.

