## Supplementary Materials for "BIQ: A method for searching circular RNAs in transcriptome databases by indexing backsplice junctions"

**Fig. 1.** Cumulative count of distinct BSJ $k$-mers in all SRA files from the (a) Drosophila and (b) ENCODE datasets.

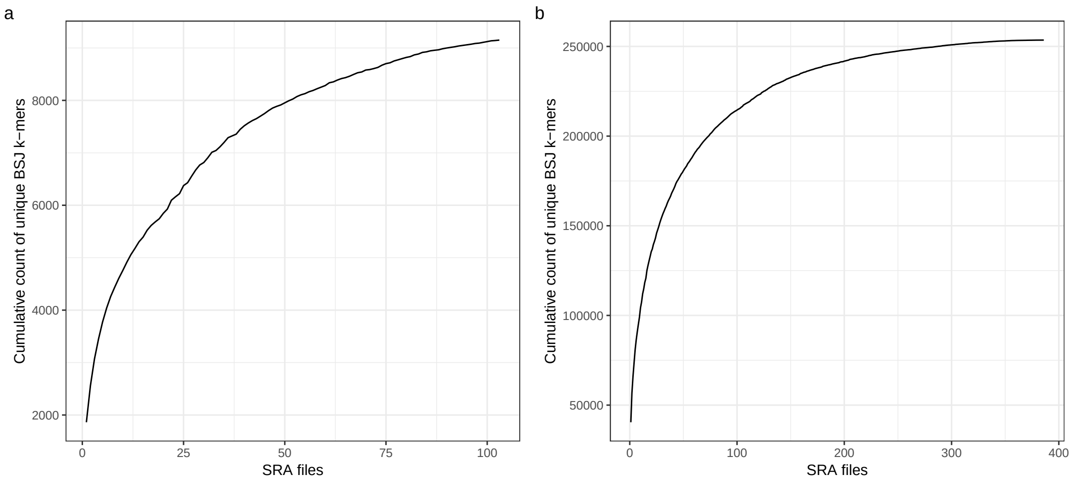


**Fig. 2.** Abundance of BSJ $k$-mers across all SRA files from the (a) Drosophila and (b) ENCODE datasets.

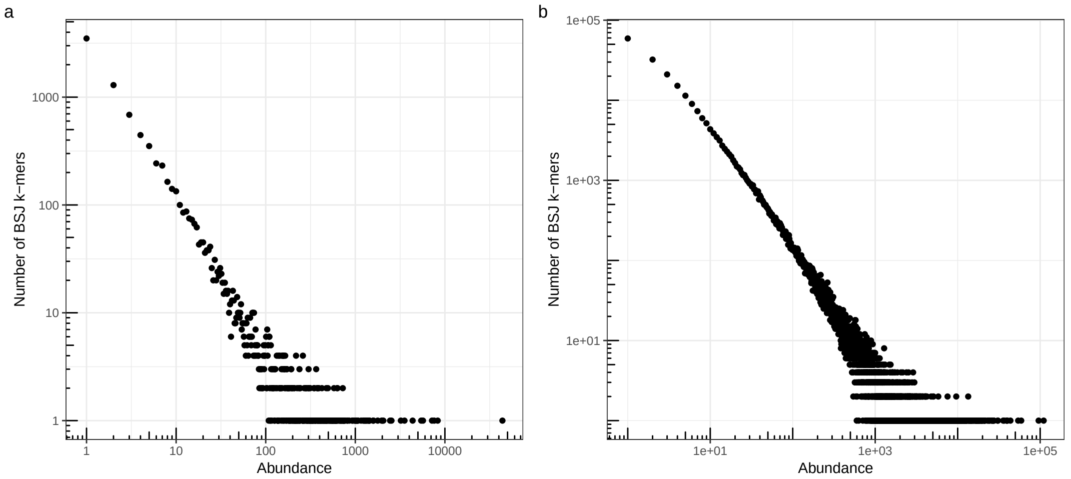


**Fig. 3.** Number of samples in which BSJ $k$-mers occur in (a) Drosophila and (b) ENCODE datasets.

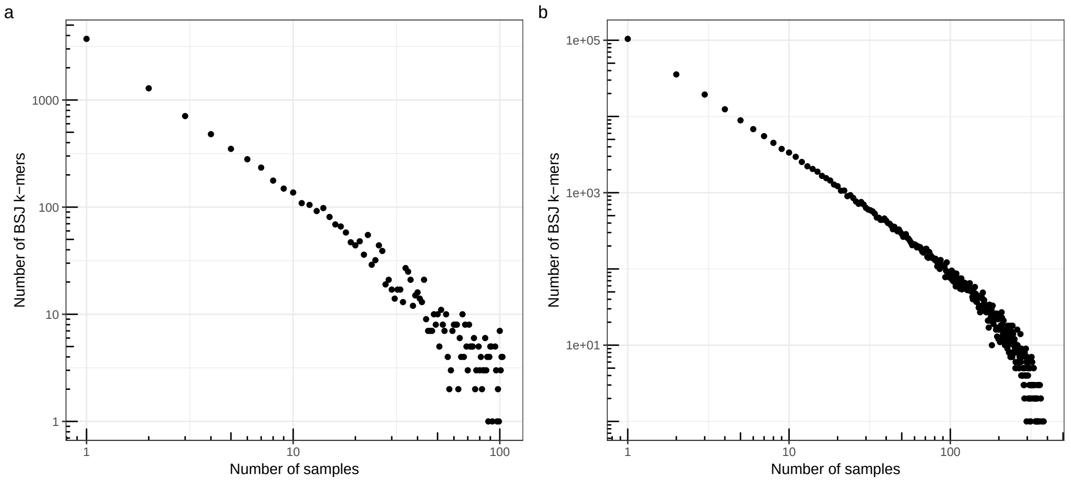

**Fig. 4.** Number of distinct observed BSJ $k$-mers (points) from *D. melanogaster* developmental transcriptomes and cumulative count of distinct BSJ $k$-mers (line)

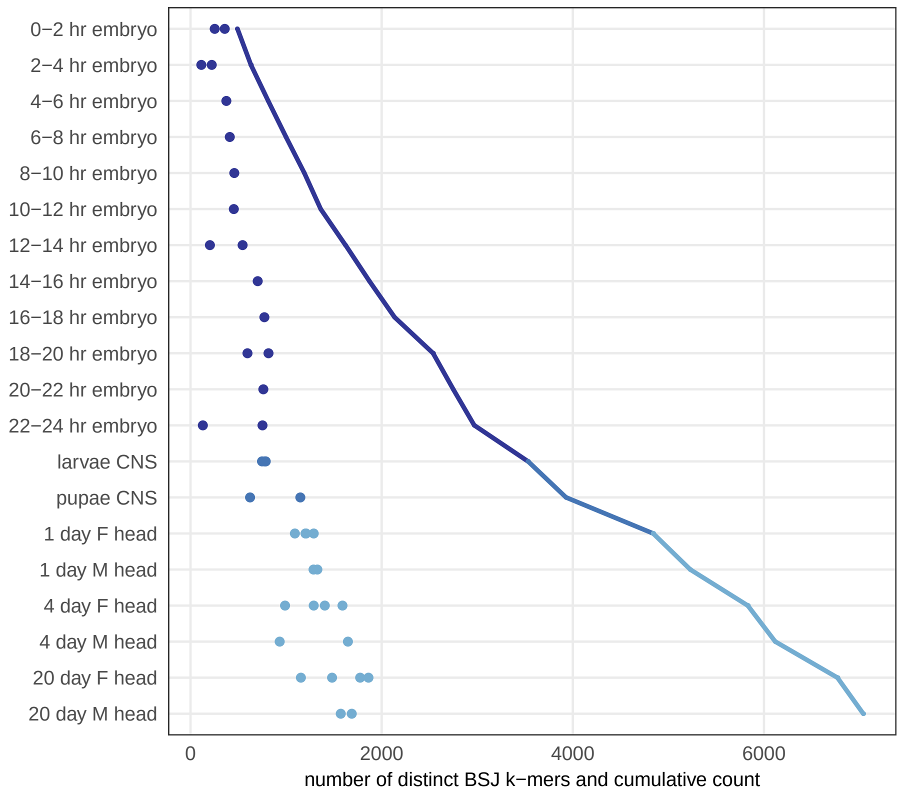

**Fig. 5.** Two-dimensional embedding of BSJ profiles of all ENCODE transcriptome datasets.

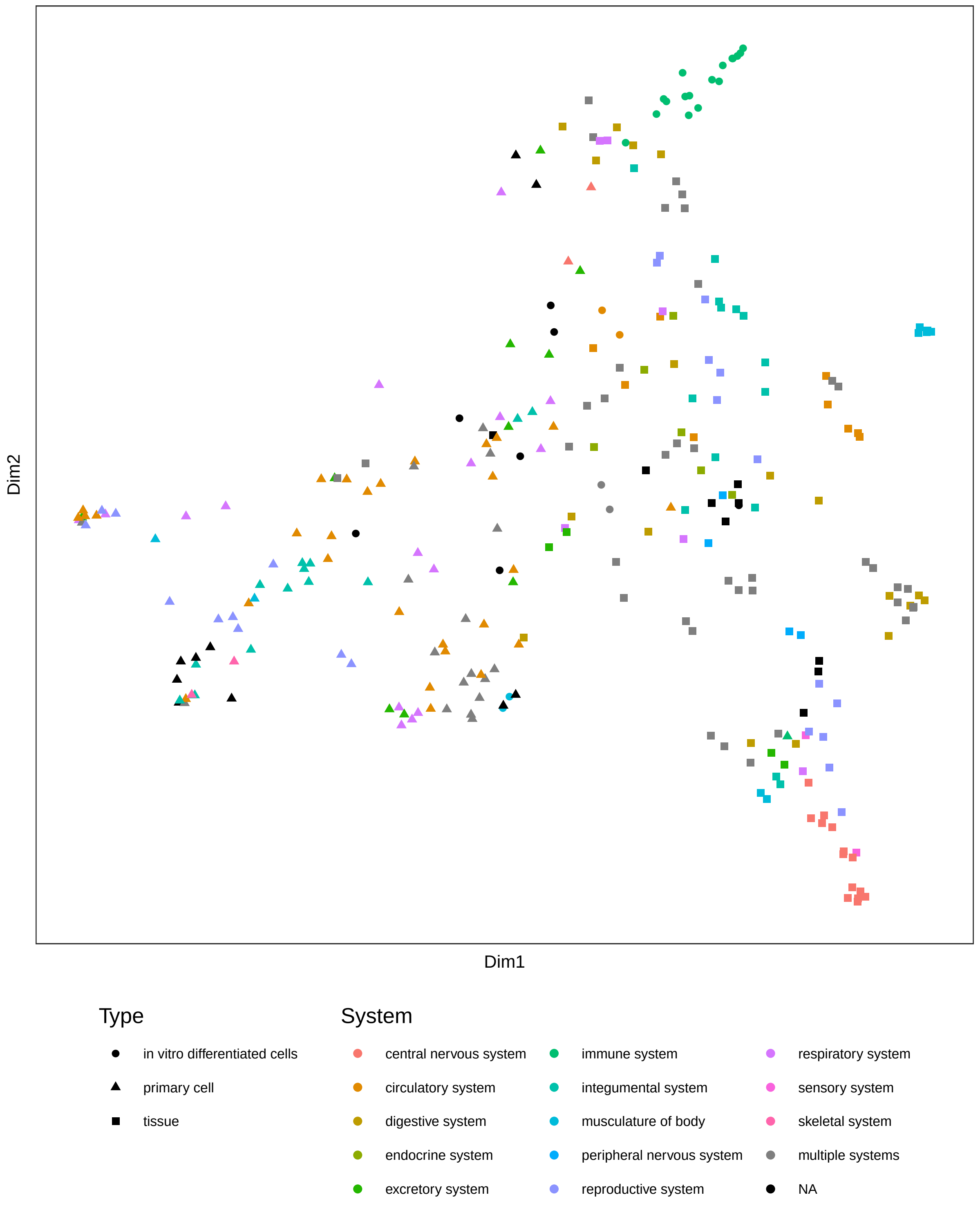
